# A robust and tunable mitotic oscillator in artificial cells

**DOI:** 10.1101/218263

**Authors:** Ye Guan, Zhengda Li, Shiyuan Wang, Patrick M. Barnes, Xuwen Liu, Haotian Xu, Minjun Jin, Allen P. Liu, Qiong Yang

## Abstract

Single-cell analysis is pivotal to deciphering complex phenomena like cellular heterogeneity, bistable switch, and oscillations, where a population ensemble cannot represent the individual behaviors. Bulk cell-free systems, despite having unique advantages of manipulation and characterization of biochemical networks, lack the essential single-cell information to understand a class of out-of-steady-state dynamics including cell cycles. Here we develop a novel artificial single-cell system by encapsulating *Xenopus* egg extracts in water-in-oil microemulsions to study mitotic dynamics. These “cells”, adjustable in sizes and periods, sustain oscillations for over 30 cycles, and function in forms from the simplest cytoplasmic-only to the more complicated ones involving nuclei dynamics, mimicking real mitotic cells. Such innate flexibility and robustness make it key to studying clock properties of tunability and stochasticity. Our result also highlights energy supply as an important regulator of cell cycles. We demonstrate a simple, powerful, and likely generalizable strategy of integrating strengths of single-cell approaches into conventional *in vitro* systems to study complex clock functions.

## INTRODUCTION

Spontaneous progression of cell cycles represents one of the most extensively studied biological oscillations. Cytoplasmic extracts predominantly from *Xenopus* eggs (Murray, 1991) have made major contributions to the initial discovery and characterization of the central cell-cycle regulators including the protein complex cyclin B1-Cdk1 (Murray et al., 1989;Lohka and Maller, 1985;Lohka et al., 1988) and the anaphase-promoting complex or cyclosome (APC/C) (Sudakin et al., 1995), as well as downstream mitotic events of spindle assembly and chromosome segregation (Hannak and Heald, 2006). Moreover, detailed dissections of the regulatory circuits in these extracts revealed architecture of interlinked positive and negative feedbacks (Kumagai and Dunphy, 1992;Mueller et al., 1995;Yang and Ferrell, 2013;Chang and Ferrell, 2013;Trunnell et al., 2011;Kim and Ferrell, 2007;Pomerening et al., 2005;Pomerening et al., 2003;Novak and Tyson, 1993b;Thron, 1996) (Figure 1A). Such interlinked feedback loops are also found in many other biological oscillators (Rust et al., 2007;Hoffmann et al., 2002;Cross, 2003;Lee et al., 2000), suggesting its importance to essential clock functions such as robustness and tunability (Tsai et al., 2008). These studies stimulated major interests in characterization of clock functions at the single cell level, for which an experimental platform is still lacking.

**Figure 1.**
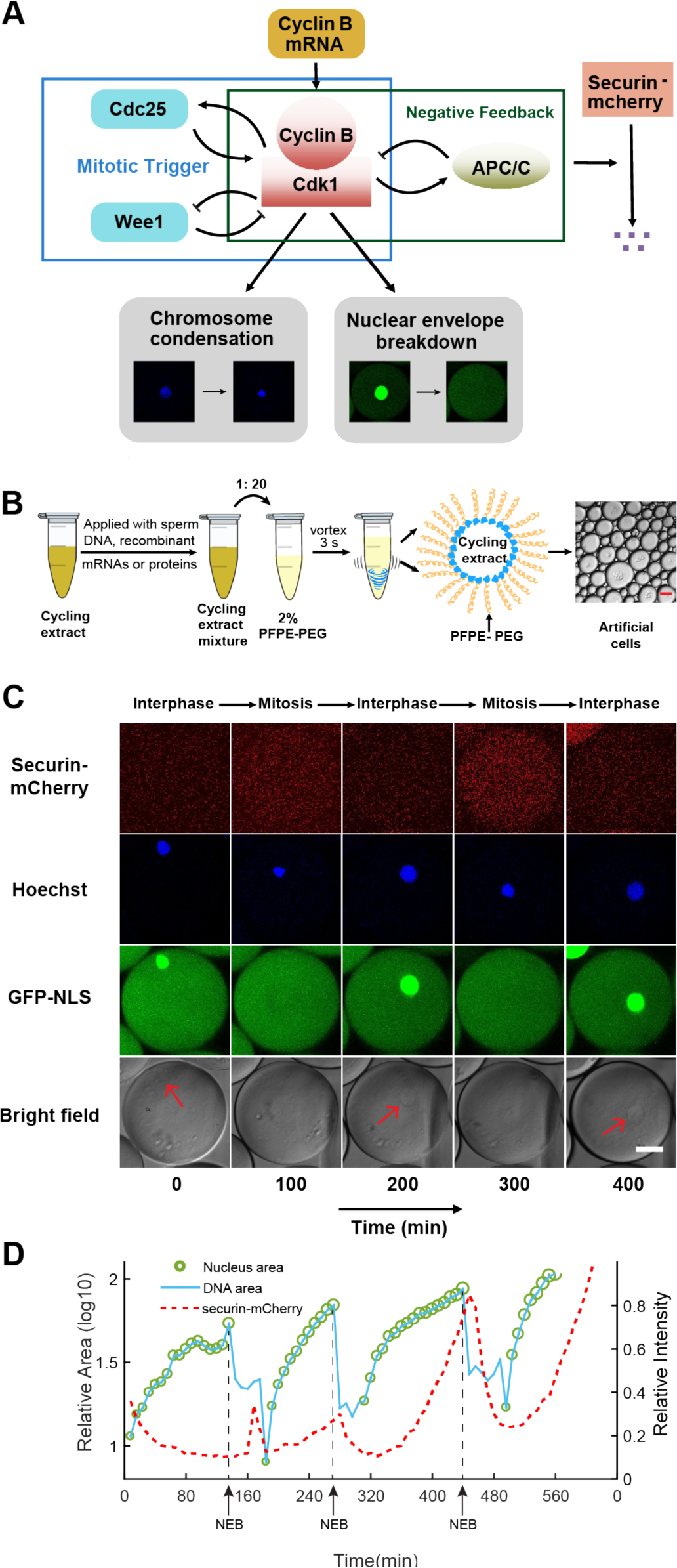
Reconstitution of an *in vitro* cell cycle clock and downstream mitotic events. A. Schematic view of a cell cycle oscillator that consists of coupled positive and negative feedback loops. The central regulator, cyclin B-Cdk1 complex activates its own activator, phosphatase Cdc25, forming a positive feedback loop, and inhibits its own inhibitor, kinase Wee1, forming a double negative feedback loop. Additionally, cyclinB-Cdk1 activates the E3 ubiquitin ligase APC/C, which targets cyclin B for degradation and completes a core negative feedback loop. Active APC/C also promotes the degradation of another substrate securin. Once cyclinB1-Cdk1 complex is activated, the circuit drives a set of mitotic events including chromosome condensation and nuclear envelope breakdown (NEB). B. Experimental procedures. Cycling *Xenopus* extracts are supplemented with various combinations of recombinant proteins, mRNAs, and de-membraned sperm DNAs, which are encapsulated in 2% Perfluoropolyether-poly (ethylene glycol) (PFPE-PEG) oil microemulsions. Scale bar is 100 μm. C. Snapshots of a droplet were taken periodically both in fluorescence channels (top three rows) and bright-field (the last row). The cyclic progression of the cell cycle clock and its downstream mitotic processes is simultaneously tracked by multiple fluorescence reporters. The clock regulator APC/C activity is reported by its substrate securin-mCherry, chromosomal morphology changes by the Hoechst stains, and NEB by GFP-NLS. Nuclear envelopes (red arrows) are also detectable on bright field images, matching the localization of GFP-NLS indicated nuclei. Scale bar is 30 μm. D. Multi-channel measurements for the droplet in Figure 1C. The nucleus area (green circle) is calculated from the area of the nuclear envelope indicated by GFP-NLS, noting that the areas of the green circles are also scaled with the real areas calculated for the nuclei. DNA area curve (blue line) shows the chromosome area identified by Hoechst 33342 dye. Chromosome condensation happens almost at the same time as the nuclear envelope breaks down (black dashed line). The red dashed line represents the intensity of securin-mCherry over time, suggesting that degradation of the APC/C substrate lags behind NEB consistently at each cycle.

Compared to *in vivo* systems, circuits reconstituted in cell-free extracts contain well defined recombinant molecules and are more amenable to systematic design, manipulation and quantitative biochemical measurements. However, one major limitation for most *in vitro* reconstitutions up to date is that oscillations are generated in well-mixed bulk solutions, which tend to produce quickly damped oscillations (Pomerening et al., 2005). Additionally, these bulk reactions lack the similarity to the actual cell dimensions and the ability of mimicking spatial organization achieved by functional compartmentalization in real cells. These limitations make it impossible to retrieve the cellular heterogeneity to investigate important and challenging questions, such as stochasticity and tunability of an oscillator.

To overcome these challenges, we developed an artificial mitotic cycle system by encapsulating reaction mixtures containing cycling *Xenopus* egg cytoplasm (Murray, 1991) in cell-scale micro-emulsions. These droplet-based cells are stable for days and keep oscillating for dozens of cycles, offering large gains in high-throughput and long-term tracking of dynamical activities in individual droplets. In this system, we successfully reconstituted a series of mitotic events including chromosome condensation, nuclear envelope breakdown and destruction of anaphase substrates such as the proteins securin and cyclin B1. The oscillation profiles of the system such as period and number of cycles can be reliably regulated by the amount of cyclin B1 mRNAs or sizes of droplets. Additionally, we found that energy may be a critical factor for cell cycle behaviors.

## RESULTS AND DISCUSSION

### The oscillator reliably drives the periodic progression of multiple mitotic events

To create a cell-like *in vitro* mitotic system, we used a simple vortexing technique (Ho et al., 2017) to compartmentalize reactions of cycling *Xenopus* egg extracts (Murray, 1991) into oil droplets, with radius ranging from 10 μm to 300 μm (Figure 1B, Materials and Methods). The droplets were loaded on a Teflon-coated chamber and recorded using long-term time-lapse fluorescence microscopy. The fluorescence time courses of each droplet were analyzed to obtain information of period, amplitude, number of cycles, droplet size, etc.

To examine the functionality of the droplet mitotic system, we added de-membranated sperm chromatin, purified green fluorescent protein-nuclear localization signal (GFPNLS), securin-mCherry mRNA and Hoechst 33342 dyes to the cytoplasmic extracts. Figure 1C demonstrates a typical artificial mitotic cell capable of reconstructing at least three mitotic processes in parallel that alternate between interphase and mitosis. The autonomous alternation of distinct cell-cycle phases is driven by a self-sustained oscillator, the activity of which was indicated by the periodic degradation of an anaphase substrate of APC/C, securin-mCherry. In interphase, the presence of sperm chromosomal DNA, labeled by Hoechst, initiated the self-assembly of a nucleus, upon which GFP-NLS protein was imported through the nuclear pores. The spatial distributions of Hoechst and GFPNLS thus coincided for an interphase nucleus. As the artificial cell enters mitosis, the chromosome condensed resulting a tighter distribution of Hoechst, while the nuclear envelope broke down and GFP-NLS quickly dispersed into a uniform distribution in the whole droplet. The time courses for these processes were analyzed in Figure 1D, indicating that the chromosome condensation and nuclear envelop breakdown (NEB) happened before securin degradation at each cycle. All together, these experiments showed that the droplet system successfully reconstituted a cell-free mitotic oscillator of Cdk1 and APC/C that can reliably drive the periodic progression of downstream events including chromosome morphology change and nuclear envelope breakdown and reassembly, like what occurs *in vivo*.

### The oscillator is effectively tunable in frequency with cyclin B1 mRNAs

The ability to adjust frequency is an important feature for an oscillator (Tsai et al., 2008). Here, we demonstrated the present system provides an effective experimental solution to the study of tunability of the clock. To avoid any interference from the complicated nuclear dynamics, we reconstituted a minimal mitotic cycle system which, in the absence of sperm chromatin, formed no nuclei. This simple, cytoplasmic-only oscillator produced highly robust, undamped, self-sustained oscillations up to 32 cycles over 4 days (Figure 2A, B and Supplementary Video 1), significantly better than many existing synthetic oscillators. To modulate the speed of the oscillations, we supplied the system with various concentrations of purified mRNAs of full-length cyclin B1 fused to YFP (cyclin B1-YFP), which function both as a reporter of APC/C activity and as an activator of CDK1. A droplet supplied with both cyclin B1-YFP and securin-mCherry mRNAs exhibited oscillations with highly correlated signals (Figure 2C), suggesting that both are reliable reporters for the oscillator activity. With an increased concentration of cyclin B1-YFP mRNAs added to the system, we observed a decrease in the average period (Figure 2D), meaning that a higher cyclin B1 concentration tends to speed up the oscillations. However, the average number of cycles (Figure 2E) was also reduced with increased cyclin B1 concentrations, resulting in a negative correlation between the lifetime of oscillations and the amount of cyclin B1 mRNAs. The extracts will eventually arrest at a mitotic phase in the presence of high concentrations of cyclin B1.

**Figure 2.**
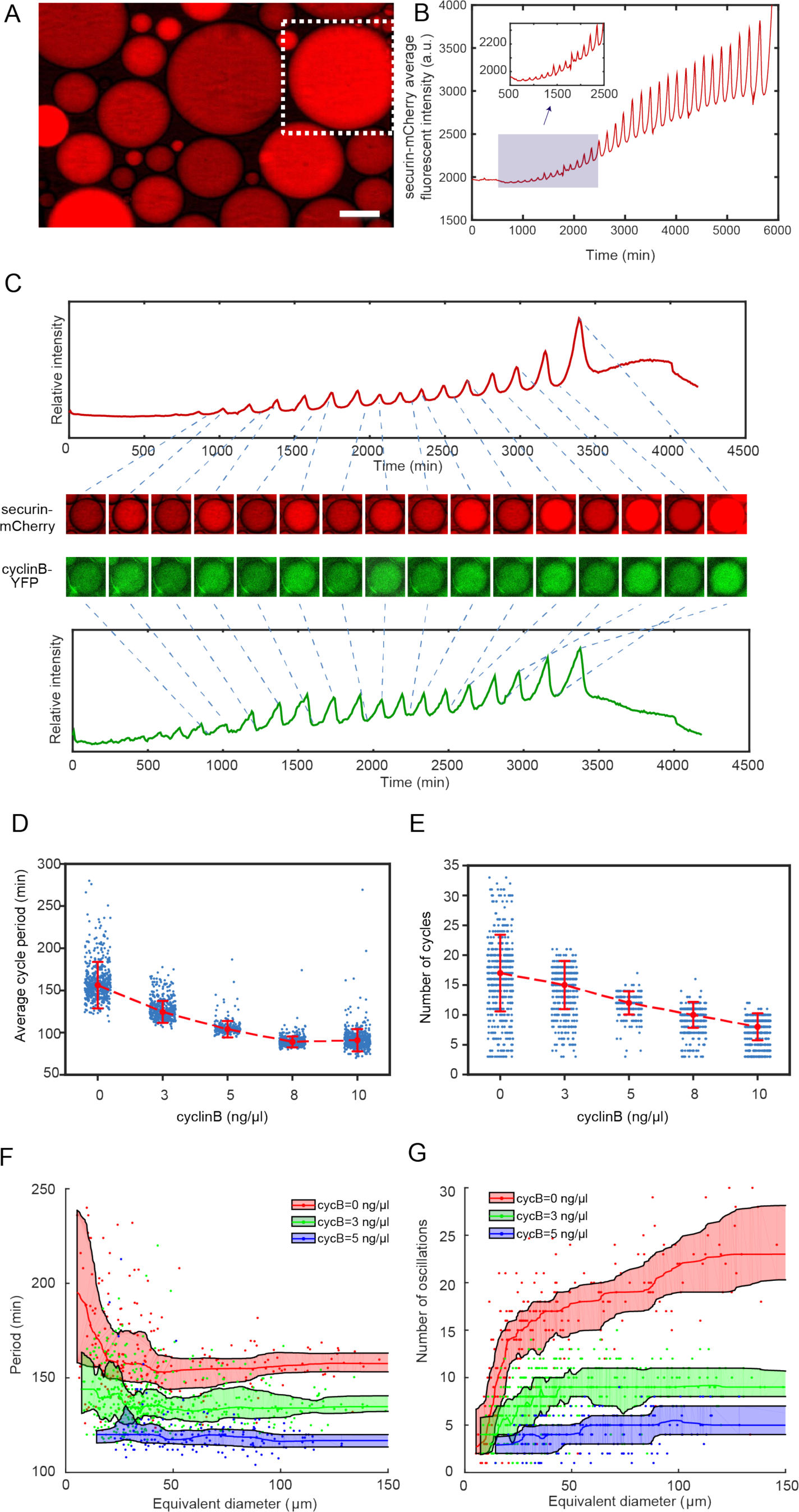
The minimal cell cycle oscillator is robust and tunable. A. Fluorescence image of securin-mCherry, a reporter for the cell cycle oscillator, in microemulsion droplets (scale bar, 100 μm). One example droplet (inside the white dotted framed square) is selected for time course analysis in Figure 2B. B. The time course of securin-mCherry fluorescence intensity of the selected droplet from Figure 2A, indicating 32 undamped oscillations over a course of 100 hours. C. Simultaneous measurements of fluorescence intensities of securin-mCherry (upper) and cyclinB-YFP (lower) within a single droplet, showing sustained oscillations for about 58 hours. The series of mCherry and YFP images correspond to selected peaks and troughs in the time courses of fluorescence intensities. The two channels have coincident peaks and troughs for all cycles, suggesting that they both are reliable reporters for the cell cycle oscillator. D, E. The oscillator is tunable in frequency (D) and number of cycles (E) as a function of the concentration of cyclin B mRNAs. Cyclin B not only functions as a substrate of APC/C but also binds to Cdk1 for its activation, functioning as an ‘input’ of the clock. In Figure 2D, the cell cycle periods are shortened by increasing the mRNA concentrations. In Figure 2E, the number of total cell cycles is reduced in response to increasing cyclin B mRNA concentrations. Red dashed line connects medians at different conditions. Error bar indicates median absolute deviation (MAD). F, G. Droplets with smaller diameters have larger periods on average and a wider distribution of periods (F), and exhibit smaller number of oscillations on average (G). Colored areas represent moving 25 percentiles to 75 percentiles with a binning size 20. The equivalent diameter is defined as the cubic root of the volume of a droplet, estimated by a volume formula in literature (Good et al., 2013). Note that these size effects are less with higher cyclin B mRNA concentrations.

### The behavior of the single droplet oscillator is size-dependent

Moreover, this system provides high flexibility in analyzing droplets with radii ranging from a few μm to above 200 μm, enabling characterization of size-dependent behaviors of cell cycles. At the scale of a cell, the dynamics of biochemical reactions may become stochastic. Although stochastic phenomenon has been studied extensively in genetic expressions, studying a system that is out of steady-state can be challenging in living organisms due to low throughput and complications from cell growth, divisions and other complex cellular environments. These limitations can be overcome by reconstitution of cell-scale *in vitro* oscillators in absence of cell growth and divisions. Parallel tracking of droplets also enables data generation for statistical analysis. Figures 2F and 2G showed that smaller droplets led to slower oscillations with a larger variance of the periods, consistent with the size effect reported on an *in vitro* transcriptional oscillator (Weitz et al., 2014). We also observed a reduced number of oscillations and a smaller variance of the cycle number in smaller droplets.

### Energy depletion model recapitulates dynamics of the oscillator

The results in Figure 2D-G indicated that the system is tunable by cyclin B1 mRNA concentration and droplet size in different manners. Although the period and number of cycles responded to varying droplet sizes in opposite directions, they followed the same trend when modulated by cyclin B1 mRNAs, resulting in a lifespan of the oscillatory system sensitive to cyclin B1 mRNA concentration. Moreover, we have observed that securin-mCherry and cyclin B1-YFP both exhibited oscillations of increased amplitude, baseline, and period over time (Figure 2C, Supplementary figure 1A, B), which cannot be explained by existing cell cycle models (Yang and Ferrell, 2013;Tsai et al., 2014).

Unlike intact embryos, cell-free extracts lack yolk as an energy source and lack sufficient mitochondria for energy regeneration. We postulated that energy is an important regulator for a droplet system with a limited amount of energy source consumed over time. To gain insights into our experimental observations and better understand the *in vitro* oscillator system, we built a model to examine how energy consumption plays a role in the oscillation behaviors. The energy depletion model is based on a well-established cell-cycle model (Yang and Ferrell, 2013;Tsai et al., 2014) modified by introducing ATP into all phosphorylation reactions (Figure 3A, Materials and methods 7 and 8).

**Figure 3.**
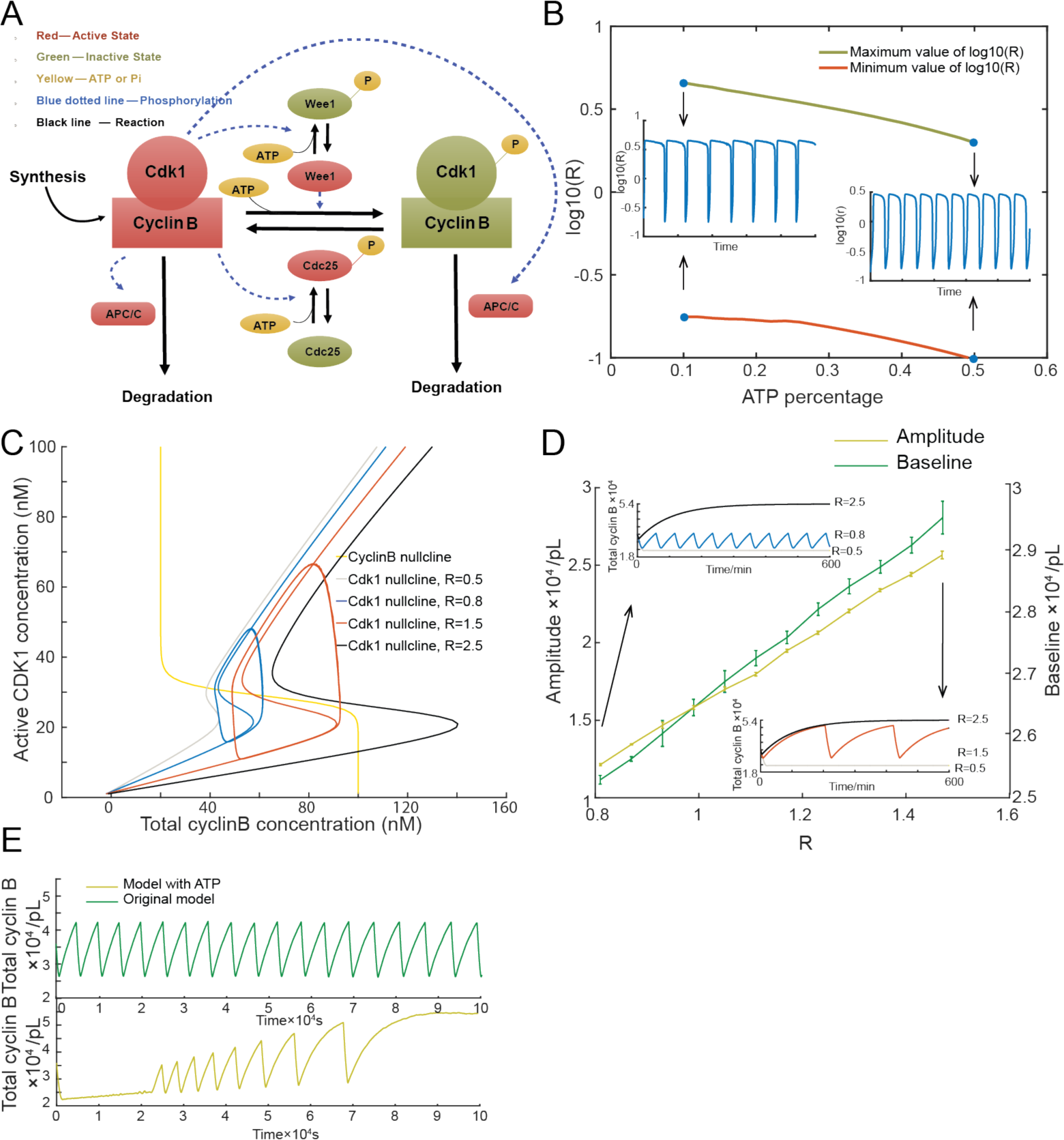
Stochastic model of cell cycle oscillations. A. Schematic view of the cyclin B-Cdk1 oscillation system. Note that ATP is taken into consideration. Activated molecules are marked in red, inactivated molecules in green and ATP or Pi in yellow. Black line indicates a reaction and blue dotted line a phosphorylation. B. Relationship between ATP percentage and *R* value (ratio of Wee1 activity to Cdc25 activity), showing that decreasing of ATP concentration leads to a higher *R* value. Two inserts represent the dynamics of R value over time when the ATP percentage [ATP]/([ATP]+[ADP]) is set as 0.1 (left) and 0.5 (right). C. Phase plots of the two-ODE model. Parameters for the cyclin B nullcline (yellow) (Yang and Ferrell, 2013), and the Cdk1 nullclines with a variety of values of *R*, were chosen based on previous experimental work (Pomerening et al., 2003;Sha et al., 2003). Two sample traces of limit cycle oscillations are plotted for R=0.8 (blue) and R=1.5 (red), showing that a larger R value leads to a higher amplitude and baseline. In addition, R=0.5 (gray) generates a low stable steady-state of cyclin B, while R=2.5 (black) a high stable steady-state of cyclin B. These stable steady-states are indicated by the intersections of the nullclines. D. Relationship between the oscillation baseline and amplitude values and ATP concentration (positively correlated with R). Error bars indicate the ranges of 3 replicates. Inserts show two example time courses of total cyclin B with different R values, colors of which match the ones in Figure 3C. E. Time series of total cyclin B molecules from the model without ATP (top panel, green line) and with ATP (bottom panel, yellow line).

In the cell cycle network, the activation of Cdk1 is co-regulated by a double positive feedback through a phosphatase Cdc25 and a double negative feedback though a kinase Wee1. The balance between Wee1 and Cdc25 activity was suggested to be crucial for the transition of cell cycle status during early embryo development (Tsai et al., 2014). In light of this, we defined the balance between Wee1 and Cdc25 by the ratio 
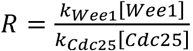
 We noted that ATP-dependent phosphorylation of Cdc25 and Wee1 can decrease *R* by activating Cdc25 and inhibiting Wee1 simultaneously, resulting in a high dependence of *R* on the ATP concentration (Figure 3B).

Using this model, we further investigated the relationship between ATP and the oscillation behaviors. In Figure 3C, the phase plot of the two-ODE model shows that at a low *R* (e.g. 0.5), the system will stay in a stable steady-state with low cyclin B concentration and at a high *R* (e.g. 2.5), the oscillation will be arrested in a stable steady-state with high cyclin B concentration. At an intermediate value, increasing *R* produced oscillations of increased amplitude, baseline and period (Figure 3C, D). If we assume that the available ATP concentration decreases over time, we can readily recapitulate the experimentally observed increment of amplitude, baseline, and period of the cyclin B time course (Figure 3E).

We noted that, besides phosphorylation, other processes, including protein synthesis and ubiquitination-mediated degradation, also consume ATPs and are sensitive to the energy level. However, the changes of synthesis and degradation rates yielded no obvious effects on the amplitude and baseline (Supplementary figure 1D).

The energy depletion model also predicted the experimental observations in Figure 2D-G by showing that increasing cyclin B concentrations decreased both period and number of cycles (Supplementary Figure 1C), while when the droplet diameter increased, the mean (and standard deviation) period decreased with increased mean (and standard deviation) number of cycles (Supplementary Figure 1E).

We have developed here a novel artificial cell system that enables highly robust and tunable mitotic oscillations. The system is amenable to high throughput, quantitative manipulation and analysis of both cytoplasmic and nuclear processes. Given cell cycles share common topologies with many biological oscillators, the system may be valuable to investigate fundamental principles of oscillator theory.

Our energy depletion model suggested an interesting mechanism to modulate oscillations with a single control parameter *R* that depends on the energy-tunable balance of two positive feedback loops. Considering that the rapid, non-stopping cell divisions of an early embryo require a large amount of energy, this energy-dependent control may function as a “checkpoint” to arrest cell cycles if *R* becomes too large.

## MATERIALS AND METHODS

### 1. Cycling Xenopus laevis extract preparation

Cycling *Xenopus* extracts were prepared as described (Murray, 1991), except that eggs were activated with calcium ionophore A23187 (200 ng/μL) rather than electric shock. Freshly prepared extracts were kept on ice while applied with de-membranated sperm chromatin (to approximately 250 per μl of extract), GFP-NLS (10 μM) and recombinant mRNAs of securin-mCherry (10 ng/μL) and cyclin B1-YFP (ranging from 0 to 10 ng/μL). The extracts were mixed with surfactant oil 2% PFPE-PEG to generate droplets.

### 2. Fluorescence-labeled reporters

GFP-NLS protein was expressed in BL21 (DE3)-T-1 competent cells (Sigma Aldrich, B2935) that were induced by 0.1 mM IPTG (Isopropyl β-D-1-thiogalactopyranoside, Sigma Aldrich, I6758) overnight. Cells were broken down to release protein through sonication. GE Healthcare Glutathione Sepharose 4B beads (Sigma Aldrich, GE17-0756-01) and PD-10 column (Sigma Aldrich, GE17-0851-01) were used to purify and elute GFP-NLS protein. 200 mg/ml Hoechst 33342 (Sigma Aldrich, B2261) was added to stain chromosomes. Securin-mCherry and cyclin B1-YFP plasmids were constructed using Gibson assembly method (Gibson et al., 2009). All mRNAs were *in vitro* transcribed and purified using mMESSAGE mMACHINE SP6 Transcription Kit (Ambion, AM1340).

### 3. Teflon-coated microchamber preparation

VitroCom miniature hollow glass tubing with height of 100 μm (VitroCom, 5012) into pieces was cut into pieces with lengths of 3-5 mm. A heating block was heated up to 95℃ in a Fisher Scientific Isotemp digital incubator and then it was placed into a Bel-art F42025-0000 polycarbonate vacuum desiccator with white polypropylene bottom. The cut glass tubes and a 1.5 ml Eppendorf tube containing 30 μl Trichloro (1H,1H,2H,2Hperfluorooctyl) silane (Sigma Aldrich, 448931) were placed in the heating block. Vacuum was applied to the desiccator and the tubes was left incubated overnight.

### 4. Generation of droplet-based artificial cells

To generate droplets, we used a Fisher Scientific vortex mixer to mix 20 μl cycling extract reaction mix, and 200 μl 2% PFPE-PEG surfactant (Ran Biotechnologies, 008FluoroSurfactant-2wtH-50G) at speed level 10 for 3 seconds. By adjusting the vibration speed and ratio between aqueous and oil phase appropriately, we can obtain droplets with various sizes, ranging from 10 μm to 300 μm.

### 5. Time-lapse fluorescence microscopy

All imaging was conducted on an Olympus FV1200 confocal microscope under MATL mode (multiple area time lapse) and Olympus IX83 microscope equipped with a motorized x-y stage, at room temperature. Time-lapse images were recorded in brightfield and multiple fluorescence channels at a time interval of 6-9 minutes for at least 12 hours up to four days.

### 6. Image analysis and data processing

We used Imaris 8.1.2 (Bitplane Inc.) for image processing. Level-set method on brightfield images was used for droplet segmentation, and autoregressive motion algorithm was used for tracking. Tracks that had less than two oscillations were discarded. Results were then manually curated for accuracy. Means and standard deviations of fluorescence intensities as well as areas of each droplet were calculated for further analysis. The volume of a droplet was calculated using the formula proposed by a previous study (Good et al., 2013). To compensate for intensity drift over time, fluorescence intensity in droplets were normalized by average intensity of the background. For period calculation, Matlab (Mathworks Inc.) was used to detect peaks and troughs over the smoothed signal of mean intensity for cyclin B-YFP and securin-mCherry. All peaks were manually curated and edited to ensure reliability.

### 7. A two-ODE model of the embryonic cell cycle and stochastic simulations

Complicated models have been proposed to describe the embryonic cell cycle oscillation (Novak and Tyson, 1993a;Ciliberto et al., 2003;Pomerening et al., 2005;Tsai et al., 2008). However, simple two-ODE models with fewer parameters are more amenable to analysis, while still capturing the general property of the feedback loops. We described the net productions of cyclin B1 and active cyclinB-Cdk1 complex [*Cdk1_a_*] using the following two equations (Yang and Ferrell, 2013;Tsai et al., 2014):

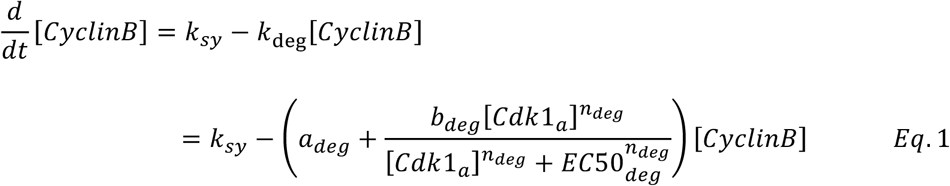

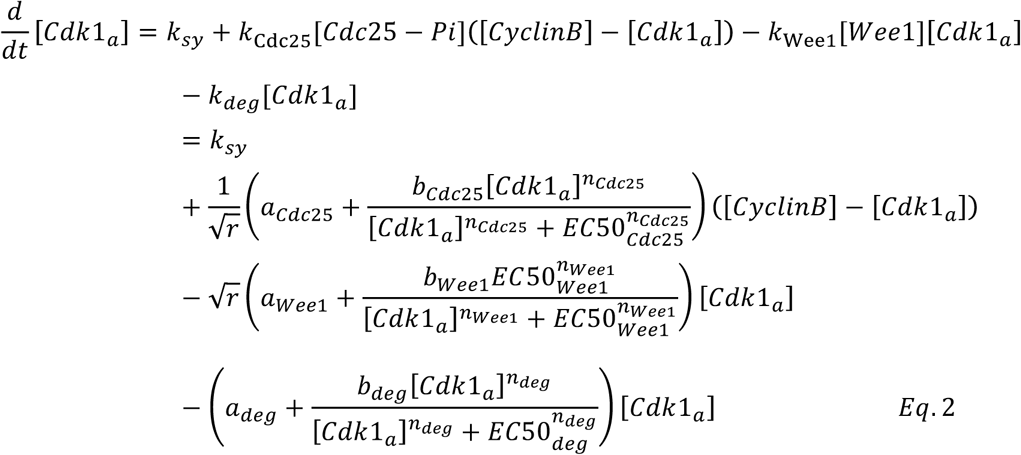

**uTable 1.**
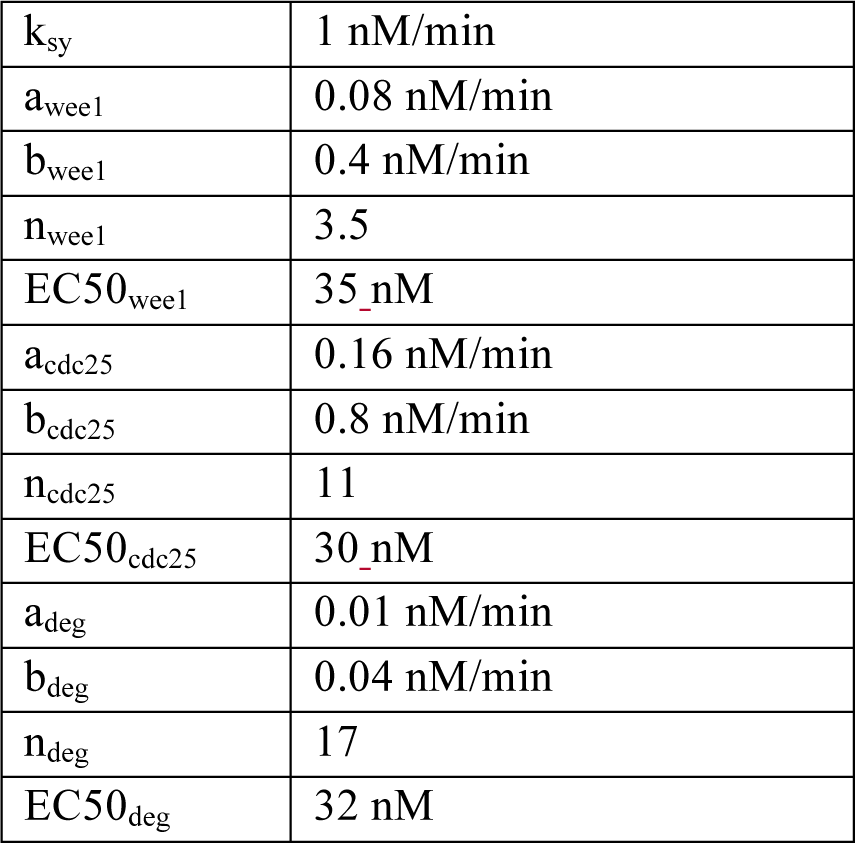
The parameters for the model are listed below:

|  |  |
| --- | --- |
| $k_{sy}$ | 1 nM/min |
| $a_{wee1}$ | 0.08 nM/min |
| $b_{wee1}$ | 0.4 nM/min |
| $n_{wee1}$ | 3.5 |
| $EC50_{wee1}$ | 35 nM |
| $a_{cdc25}$ | 0.16 nM/min |
| $b_{cdc25}$ | 0.8 nM/min |
| $n_{cdc25}$ | 11 |
| $EC50_{cdc25}$ | 30 nM |
| $a_{deg}$ | 0.01 nM/min |
| $b_{deg}$ | 0.04 nM/min |
| $n_{deg}$ | 17 |
| $EC50_{deg}$ | 32 nM |

Here, [*CyclinB*] and [*Cdk1_a_*] refer to the concentrations of cyclin B1 and active cyclin B1-Cdk1 complex. [*Wee*] is the concentration of active Wee1, while [*Cdc25 – Pi*] is the concentration of active Cdc25. We assumed that Cyclin B1 is synthesized at a constant rate. Its degradation rate is dependent on Cdk1 activity in the form of Hill function with exponent of 17 (Yang and Ferrell, 2013). Active cyclin B1-Cdk1 complex can also be eliminated through cyclin degradation. In addition, we considered that the concentration of Cdk1 to be high compared to the peak concentration of cyclin B1 (Hochegger et al., 2001;Kobayashi et al., 1991) and the affinity of these cyclins for Cdk1 to be high (Kobayashi et al., 1994). Thus, there is no unbound form of cyclin B1, and the newly synthesized cyclin B1 is converted to cyclin-Cdk1 complexes, which are rapidly phosphorylated by the Cdk-activating kinase CAK to produce active Cdk1. According to previous studies, these complexes can then be inactivated by Wee1 and re-activated by Cdc25, via the double-negative and positive feedback loops, with Hill exponent of *n_Wee1_* as 3.5 and *n_Cdc25_* as 11 (Kim and Ferrell, 2007;Trunnell et al., 2011).

We use a free parameter *r*, representing the ratio of the double negative and double positive feedback strengths, to permute the balance between the two feedbacks. This balance is suggested to be critical for oscillatory properties (Tsai et al., 2014). Note that this *r* is a parameter while R in the main text is a measurement that changes over a simulation.

In droplets that have small volumes and contain small numbers of molecules, the stochastic nature of the underlying biochemical reactions must be considered. We adapted a stochastic two-ODE model (Yang and Ferrell, 2013), and converted our two-ODE model to the corresponding chemical master equations (Kampen, 1992) and carried out numerical simulations using the Gillespie algorithm (Gillespie, 1977). The reaction rates and molecular stoichiometry are shown in Table 1.

**Table 1:**
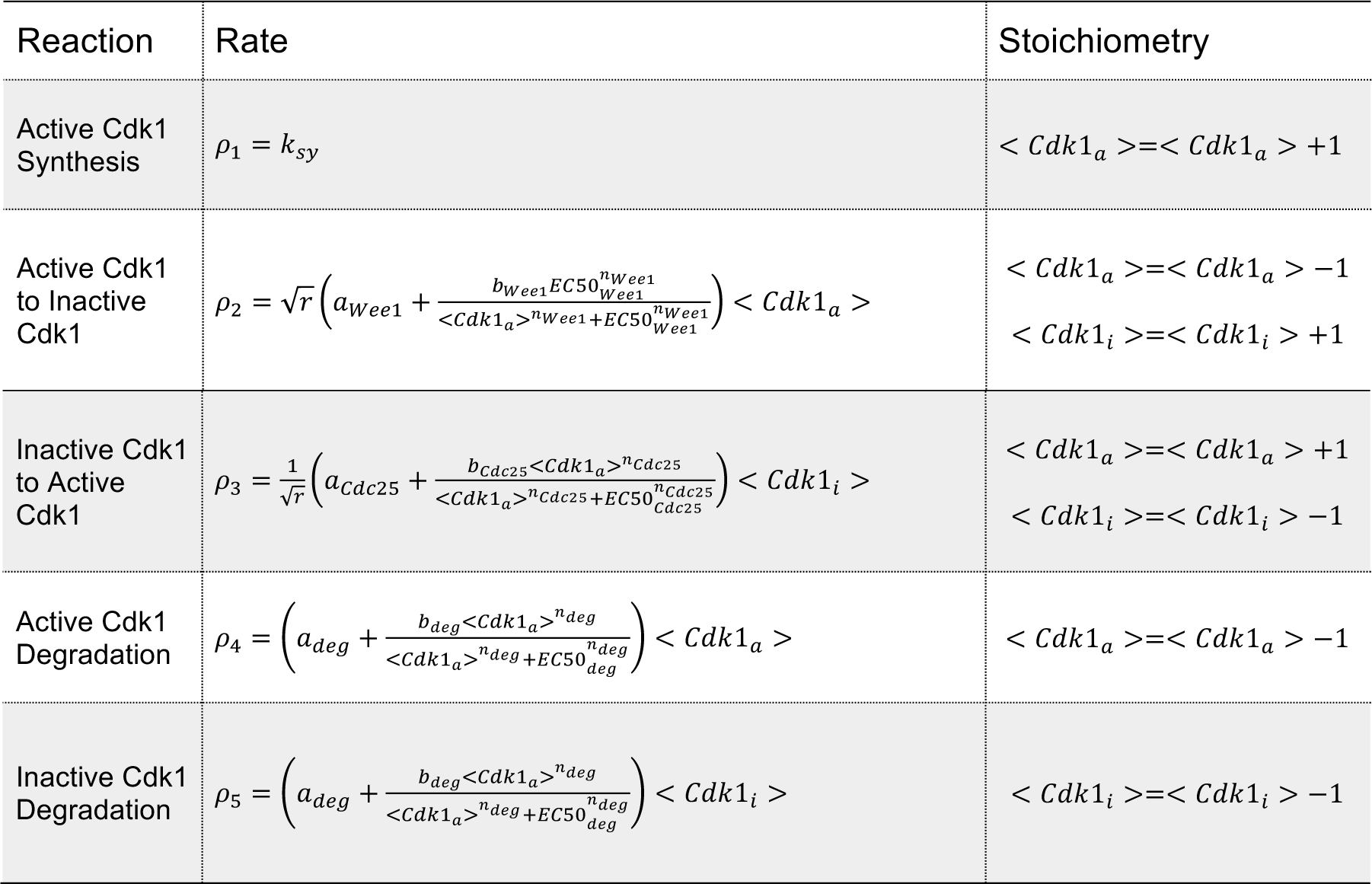
Reaction rates and stoichiometry of the two-ODE model.

| Reaction | Rate | Stoichiometry |
| --- | --- | --- |
| Active Cdk1 Synthesis | $\rho_1 = k_{sy}$ | $\langle Cdk1_a \rangle = \langle Cdk1_a \rangle + 1$ |
| Active Cdk1 to Inactive Cdk1 | $\rho_2 = \sqrt{r} \left( a_{Wee1} + \frac{b_{Wee1} EC50_{Wee1}^{n_{Wee1}}}{\langle Cdk1_a \rangle^{n_{Wee1}} + EC50_{Wee1}^{n_{Wee1}}} \right) \langle Cdk1_a \rangle$ | $\langle Cdk1_a \rangle = \langle Cdk1_a \rangle - 1$<br>$\langle Cdk1_i \rangle = \langle Cdk1_i \rangle + 1$ |
| Inactive Cdk1 to Active Cdk1 | $\rho_3 = \frac{1}{\sqrt{r}} \left( a_{cdc25} + \frac{b_{cdc25} \langle Cdk1_a \rangle^{n_{cdc25}}}{\langle Cdk1_a \rangle^{n_{cdc25}} + EC50_{cdc25}^{n_{cdc25}}} \right) \langle Cdk1_i \rangle$ | $\langle Cdk1_a \rangle = \langle Cdk1_a \rangle + 1$<br>$\langle Cdk1_i \rangle = \langle Cdk1_i \rangle - 1$ |
| Active Cdk1 Degradation | $\rho_4 = \left( a_{deg} + \frac{b_{deg} \langle Cdk1_a \rangle^{n_{deg}}}{\langle Cdk1_a \rangle^{n_{deg}} + EC50_{deg}^{n_{deg}}} \right) \langle Cdk1_a \rangle$ | $\langle Cdk1_a \rangle = \langle Cdk1_a \rangle - 1$ |
| Inactive Cdk1 Degradation | $\rho_5 = \left( a_{deg} + \frac{b_{deg} \langle Cdk1_a \rangle^{n_{deg}}}{\langle Cdk1_a \rangle^{n_{deg}} + EC50_{deg}^{n_{deg}}} \right) \langle Cdk1_i \rangle$ | $\langle Cdk1_i \rangle = \langle Cdk1_i \rangle - 1$ |

### 8. A stochastic model of the embryonic cell cycle including energy effect

To explore how energy consumption could affect the oscillations, we took ATP into account for phosphorylation and dephosphorylation of Wee1 (Tuck et al., 2013), such that:

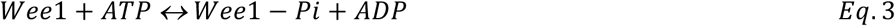

In our model, we assumed Wee1 is in equilibrium with the activity of Cdk1 due to fast reactions between Cdk1 and Wee1. Using the reaction coefficients for Wee1 phosphorylation as *k_2Wee1_* and that for Wee1-Pi dephosphorylation as *k_1Wee1_*, along with the steady-state approximation, we have:

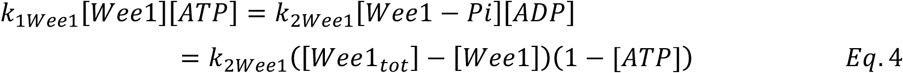

All above modifications for Wee1 reactions also applied to Cdc25. After normalizing [ATP] and [ADP] by [ATP] + [ADP], we have the updated reaction rates summarized in Table 2. Here the [wee1]_0_ and [cdc25-Pi]_0_ represent the steady-state concentration of active Wee1 and Cdc25 when ATP is not considered in reaction. The ratios of the steady-state to total concentrations of Wee1 and Cdc25 can be calculated as a function of active CDK1 using the parameters from previous work (Novak and Tyson, 1993b).

**Table 2:**
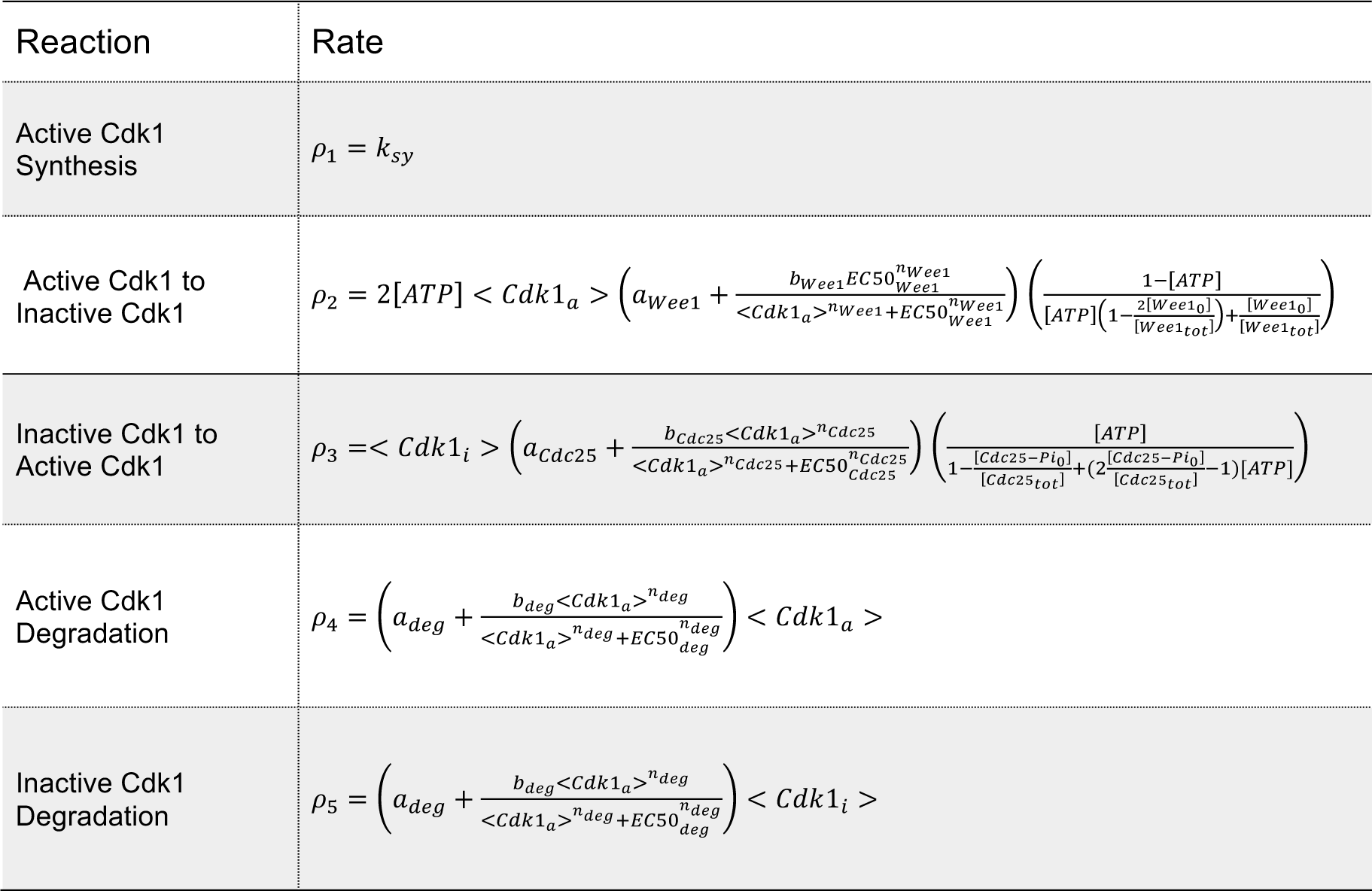
Reaction rates in the model considering ATP.

## Acknowledgements

We thank Madeleine Lu for constructing securin-mCherry plasmid, Lap Man Lee and Kenneth Ho for discussions about droplet generation, Neha Bidthanapally and Zheng Yang for helping image processing, Jeremy B. Chang and James E. Ferrell Jr for providing GFP-NLS construct. This work was supported by the National Science Foundation (Early CAREER Grant #1553031), the National Institutes of Health (MIRA #GM119688), and a Sloan Research Fellowship.

## FIGURE AND FIGURE LEGENDS

**Supplementary Figure 1.**
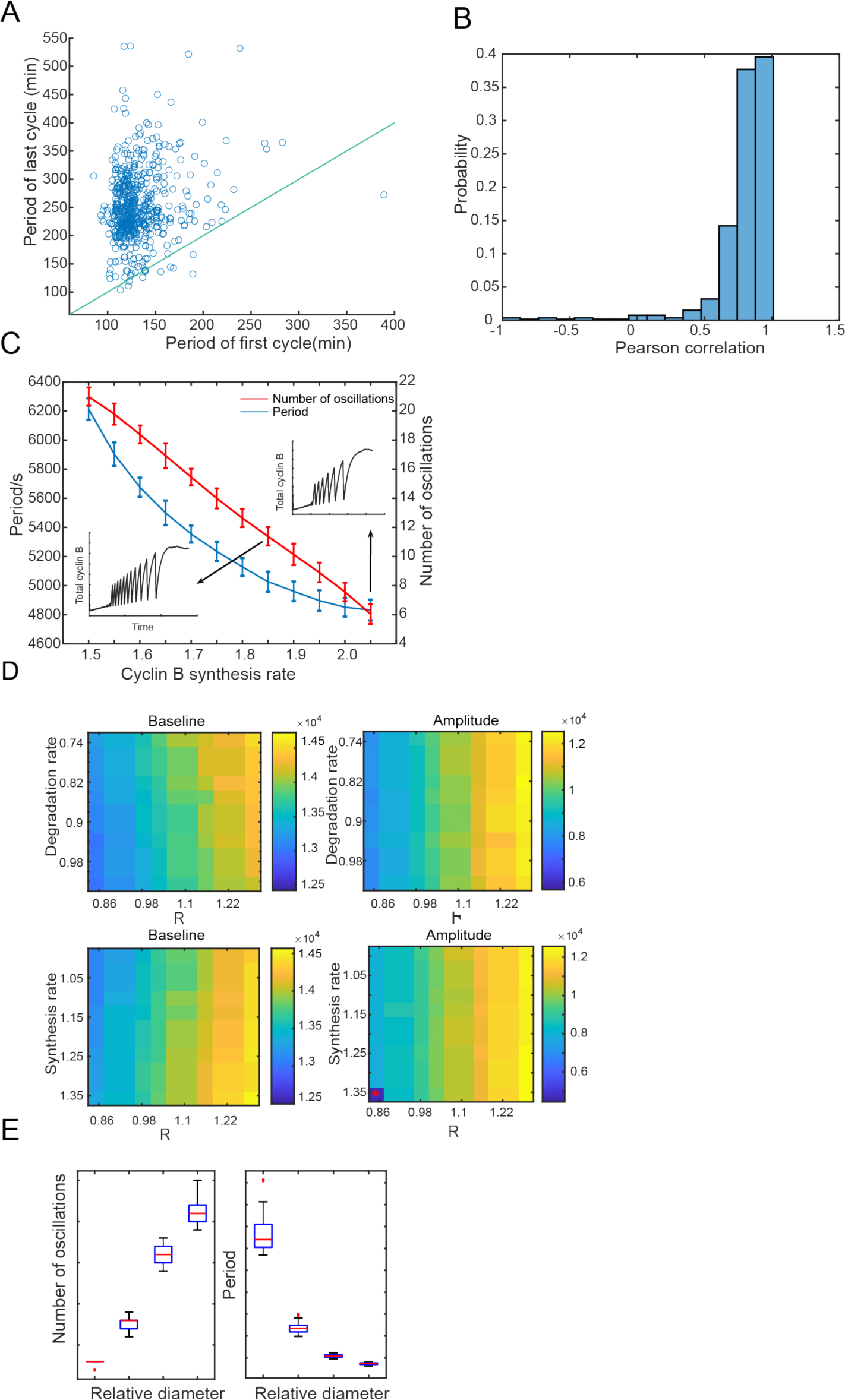
A. The period of the first cycle vs the last cycle in a droplet, showing that the period of the last cycle tends to be longer. Blue line indicates the same first and last periods. B. Pearson correlation between the period of cycle and the index of cycle in a droplet. Positive correlation indicates that the cell cycle period tends to increase over time. C. The period and number of oscillations decrease with an increasing cyclin B synthesis rate. Two inserts show example time series of total cyclin B. D. Effects of synthesis and degradation rates of cyclin B as well as R on the baseline and amplitude of oscillations of total cyclin B. Color bar indicates baseline or amplitude of cyclin B (number of molecules per pL). Asterisk means no oscillation to be observed under a certain condition. E. Effects of reaction volume on number of oscillations and period, showing that the average and variation of numbers of oscillations increase with droplet diameter and the average and variation of oscillation periods decrease with droplet diameter.

**Supplementary video 1 (Figure 1C): Reconstitution of cell cycle clock and mitotic events**

This movie corresponds to Figure 1C. Green fluorescence channel shows alternations of nuclear envelope breakdown and reformation indicated by GFP-NLS protein. Green circles disappear when nuclear envelope breakdown and reappear when nuclei assemble again. Blue channel (Hoechst) shows chromosome morphology changes over time. Securin-mCherry protein oscillations are shown in the red channel. The last channel is bright field, from which we can see nuclear envelope. Scale bar is 50 μm. The time stamp gives the real time in hour:minute format. The movie is shown at a rate of 10 frames per sec.

**Supplementary video 2 (Figure 2A, B): Free-running *in vitro* cell cycles detected by securin-mCherry reporter**

This video is from the experiment shown in Figure 2A and B. No nuclei are reconstituted in this experiment. Securin-mCherry proteins perform multiple oscillations and extract activity lasts for days. The scale bar is 100 μm and the movie is shown at a rate of 25 frames per sec.

**Supplementary video 3 (Figure 2C): Tuning of the clock speed**

This video is related to Figure 2C. The clock period can be tuned by the level of its input signal, cyclin B mRNAs. The droplets shown in the movie are applied with 3ng/μL of cyclin B1-YFP mRNAs. The YFP channel on the top shows oscillations from cyclin B1-YFP, the middle channel from securin-mCherry, and the bottom channel for the bright field. The scale bar is 100 μm and the movie is shown at a rate of 20 frames per sec.

## References Cited

Chang JB & Ferrell JE,Jr. 2013. Mitotic trigger waves and the spatial coordination of the Xenopus cell cycle. Nature, 500, 603–7. doi: 10.1038/nature12321.

Ciliberto A, Novak B & Tyson JJ 2003. Mathematical model of the morphogenesis checkpoint in budding yeast. J Cell Biol, 163, 1243–54. doi: 10.1083/jcb.200306139.

Cross FR 2003. Two redundant oscillatory mechanisms in the yeast cell cycle. Dev Cell, 4, 741–52. doi: S1534580703001199 [pii].

Gibson DG, Young L, Chuang RY, Venter JC, Hutchison CA,3rd & Smith HO 2009. Enzymatic assembly of DNA molecules up to several hundred kilobases. Nat Methods, 6, 343–5. doi: 10.1038/nmeth.1318.

Gillespie DT 1977. Exact Stochastic Simulation of Coupled Chemical-Reactions. Journal of Physical Chemistry, 81, 2340–2361.

Good MC, Vahey MD, Skandarajah A, Fletcher DA & Heald R 2013. Cytoplasmic Volume Modulates Spindle Size During Embryogenesis. Science, 342, 856–860. doi: 10.1126/science.1243147.

Hannak E & Heald R 2006. Investigating mitotic spindle assembly and function in vitro using Xenopus laevis egg extracts. Nat Protoc, 1, 2305–14. doi: 10.1038/nprot.2006.396.

Ho KK, Lee JW, Durand G, Majumder S & Liu AP 2017. Protein aggregation with poly(vinyl) alcohol surfactant reduces double emulsion-encapsulated mammalian cell-free expression. PLoS One, 12, e0174689. doi: 10.1371/journal.pone.0174689.

Hochegger H, Klotzbücher A, Kirk J, Howell M, Le Guellec K, Fletcher K, Duncan T, Sohail M & Hunt T 2001. New B-type cyclin synthesis is required between meiosis I and II during <em>Xenopus</em> oocyte maturation. Development, 128, 3795.

Hoffmann A, Levchenko A, Scott ML & Baltimore D 2002. The IkappaB-NF-kappaB signaling module: temporal control and selective gene activation. Science, 298, 1241–5. doi: 10.1126/science.1071914.

Kampen NGV 1992. Stochastic processes in physics and chemistry, Amsterdam; New York, North-Holland.

Kim SY & Ferrell JE,Jr. 2007. Substrate competition as a source of ultrasensitivity in the inactivation of Wee1. Cell, 128, 1133–45. doi: 10.1016/j.cell.2007.01.039.

Kobayashi H, Golsteyn R, Poon R, Stewart E, Gannon J, Minshull J, Smith R & Hunt T 1991. Cyclins and their partners during Xenopus oocyte maturation. Cold Spring Harb Symp Quant Biol, 56, 437–47.

Kobayashi H, Stewart E, Poon RY & Hunt T 1994. Cyclin A and cyclin B dissociate from p34cdc2 with half-times of 4 and 15 h, respectively, regardless of the phase of the cell cycle. J Biol Chem, 269, 29153–60.

Kumagai A & Dunphy WG 1992. Regulation of the cdc25 protein during the cell cycle in Xenopus extracts. Cell, 70, 139–51. doi: 0092-8674(92)90540-S[pii].

Lee K, Loros JJ & Dunlap JC 2000. Interconnected feedback loops in the Neurospora circadian system. Science, 289, 107–10.

Lohka MJ, Hayes MK & Maller JL 1988. Purification of maturation-promoting factor, an intracellular regulator of early mitotic events. Proc Natl Acad Sci U S A, 85, 3009–13.

Lohka MJ & Maller JL 1985. Induction of nuclear envelope breakdown, chromosome condensation, and spindle formation in cell-free extracts. J Cell Biol, 101, 518–23.

Mueller PR, Coleman TR & Dunphy WG 1995. Cell cycle regulation of a Xenopus Wee1like kinase. Mol Biol Cell, 6, 119–34.

Murray AW 1991. Cell cycle extracts. Methods Cell Biol, 36, 581–605.

Murray AW, Solomon MJ & Kirschner MW 1989. The role of cyclin synthesis and degradation in the control of maturation promoting factor activity. Nature, 339, 280–6. doi: 10.1038/339280a0.

Novak B & Tyson JJ 1993a. Modeling the Cell-Division Cycle - M-Phase Trigger, Oscillations, and Size Control. Journal of Theoretical Biology, 165, 101–134. doi: DOI 10.1006/jtbi.1993.1179.

Novak B & Tyson JJ 1993b. Numerical analysis of a comprehensive model of M-phase control in Xenopus oocyte extracts and intact embryos. J Cell Sci, 106 (Pt 4), 1153–68.

Pomerening JR, Kim SY & Ferrell JEJr. 2005. Systems-level dissection of the cell-cycle oscillator: bypassing positive feedback produces damped oscillations. Cell, 122, 565–78. doi: 10.1016/j.cell.2005.06.016.

Pomerening JR, Sontag ED & Ferrell JE,Jr. 2003. Building a cell cycle oscillator: hysteresis and bistability in the activation of Cdc2. Nat Cell Biol, 5, 346–51. doi: 10.1038/ncb954.

Rust MJ, Markson JS, Lane WS, Fisher DS & O’shea EK 2007. Ordered phosphorylation governs oscillation of a three-protein circadian clock. Science, 318, 809–12. doi: 10.1126/science.1148596.

Sha W, Moore J, Chen K, Lassaletta AD, Yi C-S, Tyson JJ & Sible JC 2003. Hysteresis drives cell-cycle transitions in Xenopus laevis egg extracts. Proceedings of the National Academy of Sciences, 100, 975–980. doi: 10.1073/pnas.0235349100.

Sudakin V, Ganoth D, Dahan A, Heller H, Hershko J, Luca FC, Ruderman JV & Hershko A 1995. The cyclosome, a large complex containing cyclin-selective ubiquitin ligase activity, targets cyclins for destruction at the end of mitosis. Mol Biol Cell, 6, 185–97.

Thron CD 1996. A model for a bistable biochemical trigger of mitosis. Biophys Chem, 57, 239–51.

Trunnell NB, Poon AC, Kim SY & Ferrell JE,Jr. 2011. Ultrasensitivity in the Regulation of Cdc25C by Cdk1. Mol Cell, 41, 263–74. doi: 10.1016/j.molcel.2011.01.012.

Tsai TY, Choi YS, Ma W, Pomerening JR, Tang C & Ferrell JE,Jr. 2008. Robust, tunable biological oscillations from interlinked positive and negative feedback loops. Science, 321, 126–9. doi: 10.1126/science.1156951.

Tsai TY, Theriot JA & Ferrell JE,Jr. 2014. Changes in oscillatory dynamics in the cell cycle of early Xenopus laevis embryos. PLoS Biol, 12, e1001788. doi: 10.1371/journal.pbio.1001788.

Tuck C, Zhang T, Potapova T, Malumbres M & Novak B 2013. Robust mitotic entry is ensured by a latching switch. Biol Open, 2, 924–31. doi: 10.1242/bio.20135199.

Weitz M, Kim J, Kapsner K, Winfree E, Franco E & Simmel FC 2014. Diversity in the dynamical behaviour of a compartmentalized programmable biochemical oscillator. Nat Chem, 6, 295–302. doi: 10.1038/nchem.1869.

Yang Q & Ferrell JE,Jr. 2013. The Cdk1-APC/C cell cycle oscillator circuit functions as a time-delayed, ultrasensitive switch. Nat Cell Biol, 15, 519–25. doi: 10.1038/ncb2737.

